# CD244 regulates both innate and adaptive immune axes in melanoma by inhibiting autophagy-mediated M1 macrophage maturation

**DOI:** 10.1101/2022.12.01.518620

**Authors:** Jeongsoo Kim, Tae-Jin Kim, Sehyun Chae, Sunghee Park, Hyojeong Ha, Yejin Park, Chul Joo Yoon, Jiyoung Kim, Jungwon Kim, Kyungtaek Im, Hyemin Lee, Kyunghye Lee, Jeongmin Kim, Daham Kim, Eunju Lee, Min Hwa Shin, Serk In Park, Inmoo Rhee, Jeewon Lee, Keun Hwa Lee, Daehee Hwang, Kyung-Mi Lee

## Abstract

Accumulating data have highlighted the role of monocytes/macrophages in immune escape by generating immunologically “cold” tumors that do not respond to immunotherapy. CD244 (SLAMF4, 2B4), a member of the signaling lymphocyte activation molecule family, is expressed on myeloid cells, but its precise role has not been elucidated. Using monocyte lineage-specific CD244-deficient (LysM-cre+/-CD244fl/fl;cKO) mice challenged with B16F10 melanoma, we report for the first time that CD244 negatively regulates tumor immunity by inhibiting the differentiation and functional maturation of CD11b^+^Ly6C^hi^F4/80^lo^ monocytes into CD11b^+^Ly6C^lo^F4/80^hi^ macrophages within the tumor microenvironment. CD244-deficient macrophages more effectively activated antigen-specific T cell responses compared to WT macrophages, thus delaying tumor growth in the B16F10 melanoma model. Moreover, combinatorial intervention of anti-PD-L1 antibodies with CD244-KO BMDM markedly improved tumor rejection compared to the anti-PD-L1 antibody alone or in combination with WT BMDM. Consistent with the murine data, transcriptome analysis of human melanoma tissue single-cell RNA-sequencing dataset (SCP398 from single-cell portal), revealed 221 differentially expressed genes of CD244^-^ monocytes/macrophages were associated with phagocytosis, antigen presentation, and autophagy. Additionally, cell type deconvolution analysis within melanoma patients bulk RNA-seq datasets from TCGA database, revealed presence of CD244- monocytes/macrophages significantly increased patient survival in primary and metastatic tumors. Hence, we proposed that CD244 serve as a critical immune checkpoint receptor on macrophages, and CD244-deficient macrophages may represent a novel therapeutic modality to convert immunologically “cold” tumors to “hot” tumors, which can function synergistically with checkpoint blockade therapies.

## INTRODUCTION

Immune exhaustion, defined by poor anti-tumor effector function with decreased proinflammatory cytokine production, contributes to an immunosuppressive tumor microenvironment (TME). The characterization of molecular pathways leading to immune exhaustion has been primarily centered on T cells, which acquire various immunomodulatory checkpoint receptors, including programmed cell death-1 (PD-1), cytotoxic T-lymphocyte-associated antigen-4 (CTLA-4), lymphocyte-activation gene-3 (LAG-3), T-cell immunoglobulin and mucin-domain containing-3 (TIM-3), and CD160, within the immunosuppressive TME ^1–4^. Accordingly, therapeutic antibodies targeting PD-1, programmed cell death ligand-1(PD-L1), and CTLA-4 have been developed to counter this immune exhaustion during the treatment of advanced cancers ^5–7^. However, treatment responses remain suboptimal for many patients due to the intrinsic and extrinsic immune evasion mechanisms, including tumor heterogeneity, lack of immune cell infiltration, and impaired antigen presentation ^8–12^.

CD244 belongs to the signaling lymphocyte activation molecule family (SLAMF) that acts as a unique immunomodulatory transmembrane receptor ^13^. Within natural killer (NK) cells and CD8 T cells, a dual co-activation and co-inhibition role for CD244 has been described ^13, 14^. Indeed, CD244 inhibitory signaling is critical for the maintenance of an exhausted NK cell and T cell phenotype within the TME ^8, 15–17^. The cytoplasmic domain of CD244 includes four immunoreceptor tyrosine-based switch motifs (ITSMs) that interact with at least five distinct adaptor molecules: SAP, EAT-2, SHP1, SHP2, and SHIP-1. Intracellular binding of SAP propagates an activating signal, whereas binding of SH2 phosphatases propagates an inhibitory signal. Hence, whether CD244 propagates an activating or inhibitory signal is determined by the expression levels, availability, and competitive binding of CD244 and adaptor molecules^18^.

Monocytes are indispensable for the initiation of innate immune responses as they differentiate into macrophages within tissues and bridge the innate and adaptive immune responses through phagocytosis and antigen presentation, respectively. Accumulating data highlight the immunosuppressive phenotypes of monocytes and macrophages in the TME that suppress T cells and NK cells through nutrient depletion, inhibitory cytokine secretion, and immune checkpoint ligand expression, thus generating immunologically “cold” tumors that do not respond to immunotherapy^19–22^. More specifically, patients with advanced-stage hepatocellular carcinoma exhibit high infiltration of monocytes/macrophages into the peritumoral stroma, which is positively correlated with a severe reduction in the number of effector NK cells secreting tumor necrosis factor (TNF)-α and interferon (IFN)-γ ^15^. Furthermore, NK cells become severely exhausted upon exposure to tumor-derived monocytes expressing high levels of CD48, a CD244 ligand, in hepatocellular carcinoma tissues. Such monocyte-induced NK cell dysfunction is markedly attenuated by blocking CD244 on NK cells without impacting NKG2D or NKp30. Therefore, unidirectional signaling of CD244 bound to CD48 on monocytes drives negative signaling in NK cells.

Interestingly, CD244 is expressed not only on CD8 T and NK cells but also on multiple myeloid lineage cells, including monocytes ^23–28^. In fact, bidirectional signaling through CD244 on monocytes/macrophages may occur by interacting with neighboring CD48-expressing hematopoietic cells. Therefore, in the current study, to test this hypothesis, we crossed LysM-Cre with CD244-Flox mice to generate monocyte-specific conditional KO mice (LysM-cre^+/-^ CD244^fl/fl^; cKO) and investigated whether CD244 expression on monocytes impacts immune dysfunction and exhaustion within the TME. The murine study results were then verified in human patients through melanoma single-cell and bulk RNA sequencing datasets. The findings of this study highlight the potential immune checkpoint function of CD244 on monocytes/macrophages, while providing practical evidence for the application of CD244 blockades as therapeutic agents to convert immunologically “cold” tumors into “hot” tumors.

## RESULTS

### Absence of CD244 signaling in monocytes suppresses melanoma tumorigenesis

To determine the functional impact of CD244 expression on tumor progression, we compared tumor growth in wild-type (WT) and CD244 whole-body KO (wKO) mice subcutaneously injected with 1 × 10^6^ B16F10 tumor cells. The growth of B16F10 tumors in CD244 wKO mice was significantly lower than that in WT mice (Fig. 1A), similar to head and neck squamous cell carcinoma model reported previously ^29^. When analyzed 14 days after tumor inoculation, the proportion of lymphoid (CD4 T, CD8 T, and NK cells) and myeloid (monocytes, macrophages, neutrophils, dendritic cell [DC]) cells in the spleen and tumor was comparable in WT and CD244 wKO mice, as assessed by flow cytometry (Supplementary Fig. 1A). Strikingly, CD244 expression was specifically upregulated in CD45+ leukocytes harvested from the tumor but not the spleen (Fig. 1B and C). When analyzed the proportion of immune cell subtypes expressing CD244 among the total CD45+ cells in the tumor, significant upregulation of CD244 was noted in monocytes (39.1-fold) and macrophages (41.2-fold; Fig. 1D), but not in neutrophils (9.6-fold), compared to the proportion in spleen.

**Fig. 1.**
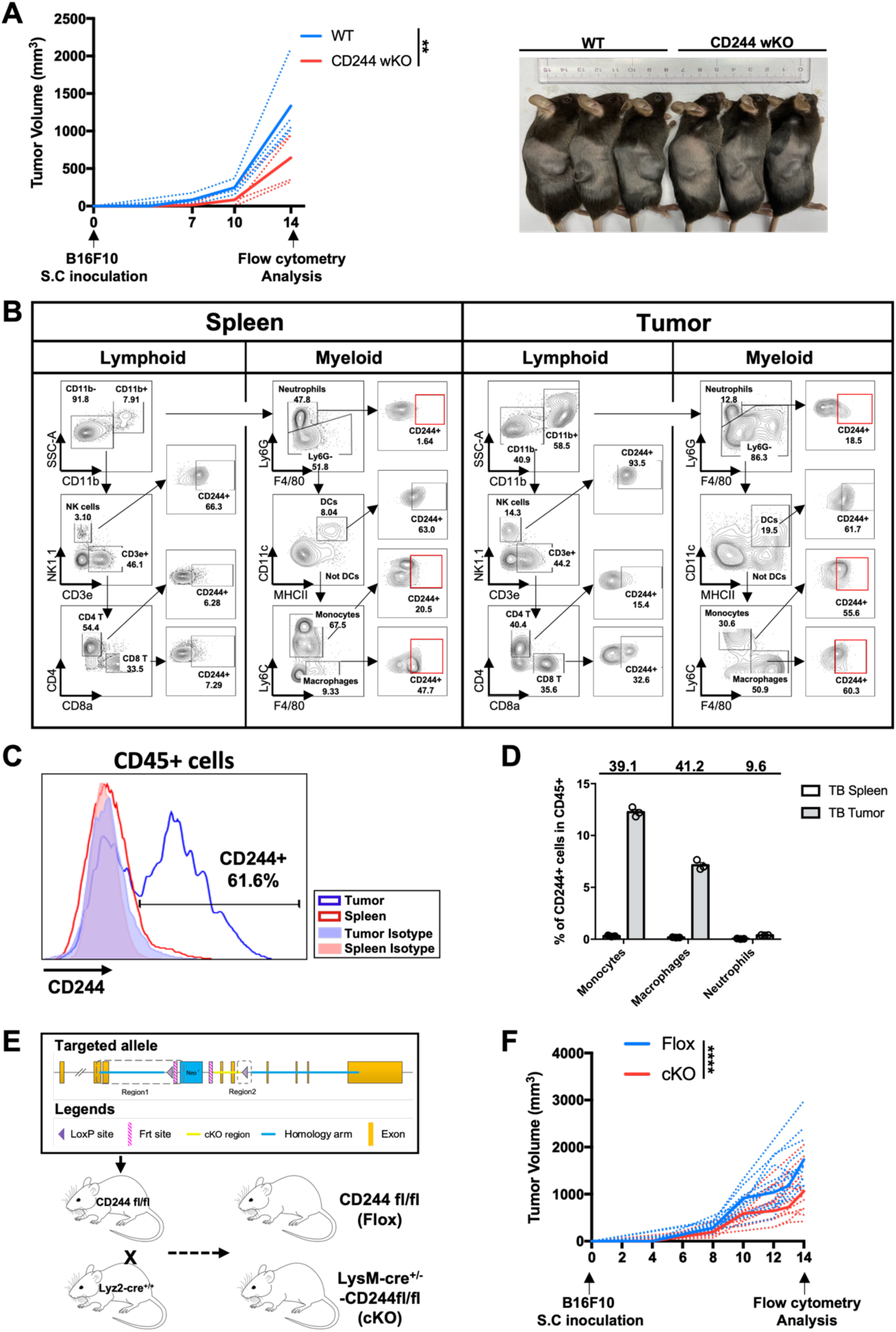
Monocytes/macrophages expressing CD244 are increased in tumor micro- environment and accelerate tumor growth. (**A**) **(left)** B16F10 tumor growth in C57BL/6(WT) and CD244^-/-^ (CD244 wKO) mice (n = 4, N = 3; two-way ANOVA). Representative photograph of WT and CD244 wKO mice 14 days after B16F10 inoculation **(right)**. (**B**) CD244 expression on lymphoid and myeloid cells was measured by flow cytometry in spleen **(left)** and tumor **(right)** (n = 10). (**C**) Representative histogram of CD244 expression levels on whole CD45+ cells in WT spleen and tumor (n = 7). (**D**) Proportion of CD244-expressing myeloid cell subpopulations among CD45+ cells in spleen and tumor of tumor bearing mice (n = 3). Data is presented as fold change in proportion of CD244 expressing cells between the spleen and tumor mass of Tumor bearing (TB) mice. (**E**) Scheme for generating littermate CD244^fl/fl^ (Flox) and LysMcre^-/+^ CD244^fl/fl^ (cKO) mice. (**F**) B16F10 tumor growth in CD244^fl/fl^ (Flox) and LysMcre^-/+^ CD244^fl/fl^ (cKO) mice (n = 13, N = 3; two-way ANOVA).

We next investigated whether elevated CD244 expression on monocytes and macrophages within tumors is associated with tumor immune escape, as previously observed with NK cells and CD8 T cells in human patients and mouse models ^15, 17, 29–32^. To this end, we generated C57BL/6 background CD244^fl/fl^ (Flox) mice and crossed them with LysM-cre^+/-^ mice to produce LysM-cre^+/-^ CD244^fl/fl^ (cKO) mice, in which CD244 was deleted specifically in the monocyte lineage cells (Fig. 1E and Supplementary Fig. 1B). These mice were then injected subcutaneously with syngeneic B16F10 melanoma cells (Supplementary Fig. 1C). Similar to the results shown for CD244 whole KO mice (wKO), monocyte-specific deletion of CD244 in cKO mice resulted in a significant reduction in tumor growth compared with the growth in littermate control Flox mice (Fig. 1F). Together, these data demonstrate the previously unknown function of CD244 on monocytes as an important negative regulator of tumor immunity.

### Loss of CD244 on CD11b+Ly6C^hi^ monocytes alters tumor microenvironment, favoring differentiation into M1 macrophages

To delineate the mechanism underlying accelerated tumor clearance in cKO mice, we first examined their myeloid cell populations within the tumor mass in comparison to those in control Flox mice. No significant differences in the proportion of CD45^+^CD11b^+^ myeloid cells were observed between Flox and cKO mice (Fig. 2A, left). Furthermore, neither the proportion of CD11b^+^Ly6G^+^ neutrophils (Fig. 2A, middle) nor CD11c^+^MHCII^+^ DCs (Fig. 2A, right) was altered in CD244 cKO mice. However, the proportion of CD11b^+^Ly6G^-^Ly6C^hi^F4/80^lo^ monocytes was significantly decreased, while that of CD11b^+^Ly6G^-^Ly6C^lo^F4/80^hi^ macrophages was significantly increased in cKO mice compared to those in Flox mice (Fig. 2B). In contrast, the proportion of CD206-expressing M2 macrophages was comparable between CD244 cKO and Flox mice (Fig. 2C). Increased macrophages in cKO mice were accompanied by significant upregulation of M1 markers and cytokines, including inducible nitric oxide synthase (iNOS), interleukin (IL)-6, and interferon (IFN)-β, in CD11b^+^ tumor-infiltrating myeloid cells, while the expression of arginase-1 (ARG-1) and transforming growth factor (TGF)-β, which represent M2 macrophages, was not altered (Fig. 2D). These results prompted us to investigate whether the absence of CD244 in the monocyte lineage impacts monocyte differentiation into M1 macrophages.

**Fig. 2.**
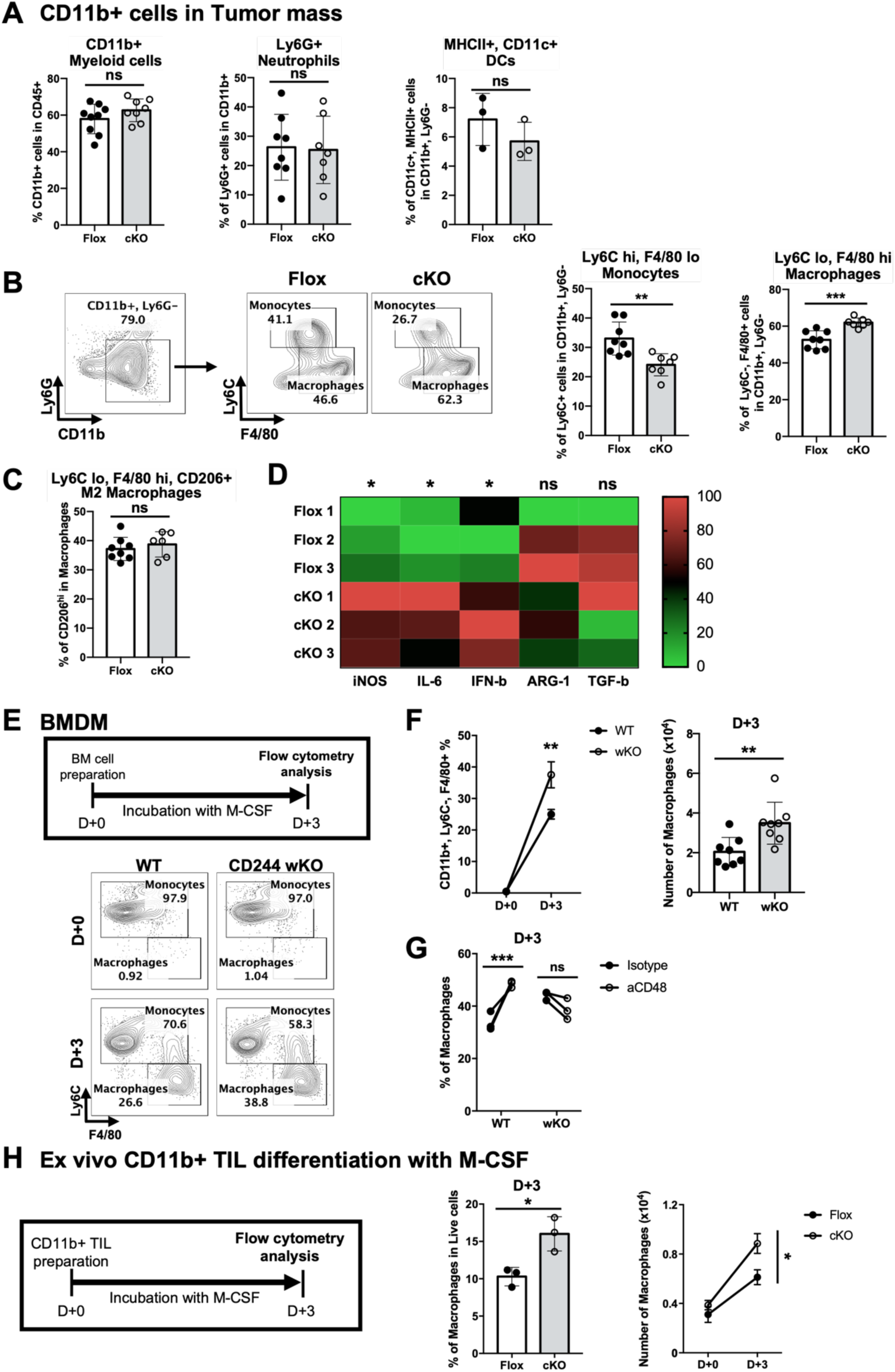
Absence of CD244 enhances M1 macrophage differentiation. (**A–C**) Proportion of myeloid populations in tumor mass of Flox (n = 8) and cKO (n = 6 or 7) mice at 14 days following B16F10 inoculation. N=2; unpaired Student’s *t*-test. Proportion of (**A**) CD11b+ myeloid cells in CD45+ cells **(left)**, CD45+, CD11b+ Neutrophils **(center)**, and CD45+, CD11b+, Ly6G- DCs **(right)**. (**B**) Gating strategies **(left)** and proportion of monocytes (Ly6C^high^, F4/80^low^) and macrophages (Ly6C^low^, F4/80^high^) in CD45+, CD11b+, Ly6G- cells **(right)**. (**C**) Proportion of M2 macrophages (Ly6C^low^, F4/80^high^, CD206^+^) in CD11b^+^, Ly6G^-^, Ly6C^low^, F4/80^high^ macrophages. (**D**) mRNA expression of M1 and M2 macrophage genes in CD11b+ tumor-infiltrating leukocytes. n = 3; unpaired Student’s *t*-test. (**E**) Experiment scheme and representative flow cytometry plot of bone marrow-derived macrophages (BMDM) from CD244 wKO and WT mice. (**F**) Proportion of WT and CD244 wKO macrophages among CD45+ cells on D+0 and D+3 **(left)**; n = 3, N = 3; two-way ANOVA. Absolute macrophage numbers **(right)**. n = 8, N = 3; unpaired Student’s *t*- test. (**G**) Macrophage proportion in WT and CD244 wKO BMDM cultures treated with isotype or anti-CD48 antibody. n = 3, N = 2; paired *t*-test. (**H**) Experiment scheme **(left)**, proportion **(center),** and abundance of **(right)** macrophages in tumor infiltrating CD11b+ cells cultured with M-CSF. n = 3; unpaired Student’s *t*-test **(left)** and two-way ANOVA **(right)**.

Accordingly, we set up an *in vitro* macrophage differentiation assay using whole bone marrow (BM) cells in the presence of 50 ng/mL recombinant macrophage colony-stimulating factor (M-CSF; Fig. 2E). Results revealed a significant increase in the proportion (Fig. 2F, left) and number (Fig. 2F, right) of CD11b^+^Ly6C^lo^F4/80^hi^ macrophages in CD244 wKO bone marrow- derived macrophages (BMDM) (37.5% ± 0.030, 3.5 × 10^4^ ± 0.37 × 10^3^) in comparison to those in WT BMDM (25.0% ± 0.021, 2.0 × 10^4^ ± 0.26 × 10^3^). To further investigate whether the increased macrophage population in CD244-deficient mice was associated with defective signaling downstream of ligation with CD48, we differentiated BM monocytes in the presence of anti-CD48 monoclonal antibodies (mAbs) to block their interactions with CD244. Compared to the isotype control, anti-CD48 mAb treatment significantly increased the macrophage population in the WT culture, but not in the CD244 wKO culture (Fig. 2G). These data suggest that CD48 ligation with CD244 is critical for suppressing differentiation of M1 macrophages.

To exclude the possibility that the physiological properties of BMDM and tumor-located monocytes are significantly different, tumor-infiltrating CD11b+ cells from cKO and Flox mice were isolated and subjected to an *in vitro* macrophage differentiation assay. Tumor- infiltrating myeloid cells were incubated with M-CSF; the proportion and number of CD11b^+^Ly6C^lo^F4/80^hi^ macrophages was then assessed 3 days after culture. Similar to the results with whole BM cells, tumor-infiltrating monocytes isolated from CD244 cKO mice showed a significantly increased proportion and number of macrophages compared to those obtained from Flox mice (Fig. 2H). Taken together, the specific loss of CD244 expression in CD11b^+^Ly6C^hi^F4/80^lo^ monocytes favors differentiation into CD11b^+^Ly6C^lo^F4/80^hi^ macrophages within the tumor mass.

### Targeted CD244 deletion in monocyte-lineage cells enhances antigen-specific anti-tumor immunity

To further investigate the underlying mechanisms of tumor regression by monocyte-specific deletion of CD244, we harvested tumor infiltrating lymphocyte (TIL)s from B16F10 tumor-bearing Flox and cKO mice and quantified IFN-γ expression by flow cytometry in response to irradiated B16F10 tumor targets *ex vivo*. Although the proportion of lymphocytes was not altered (Fig. 3A), IFN-γ expression in CD8 T cells (Fig. 3B, upper), and CD4 T cells (Fig. 3B, lower) were significantly higher in cKO mice than in Flox mice. In addition, granzyme-b expression in CD8 T cells was significantly increased in cKO mice compared to that of Flox mice (Fig. 3C). To determine whether the increased IFN-γ and granzyme-b expression in T cells were due to an antigen-specific response, the proportion of antigen-experienced T cells expressing CD44^33^ was measured. The proportion of CD44+ CD8 T cells was significantly increased in the tumors of cKO mice compared to that in Flox mice (Fig. 3D, upper). Although the proportion of CD44+ CD4 T cells was also slightly increased in cKO mice compared to that in Flox mice, it did not show statistical significance. As expected, IFN-γ expression in CD4 T and CD8 T cells was significantly increased only in antigen-experienced CD44+ populations, not in CD44- populations (Fig. 3E and Supplementary Fig. 2A). Furthermore, the proportion of melanoma-specific H-2Db gp100 tetramer-positive CD8 T cells was significantly increased in cKO mice compared with that in Flox mice (Fig. 3F). Additionally, IFN-γ expression in CD8 T cells and CD4 T cells was significantly increased in the tumor-draining lymph nodes (TDLN) of cKO mice compared to that in Flox mice (Fig. 3G), highlighting that CD244 expressed on monocyte-lineage cells enhances antigen-specific anti-tumor immunity.

**Fig. 3.**
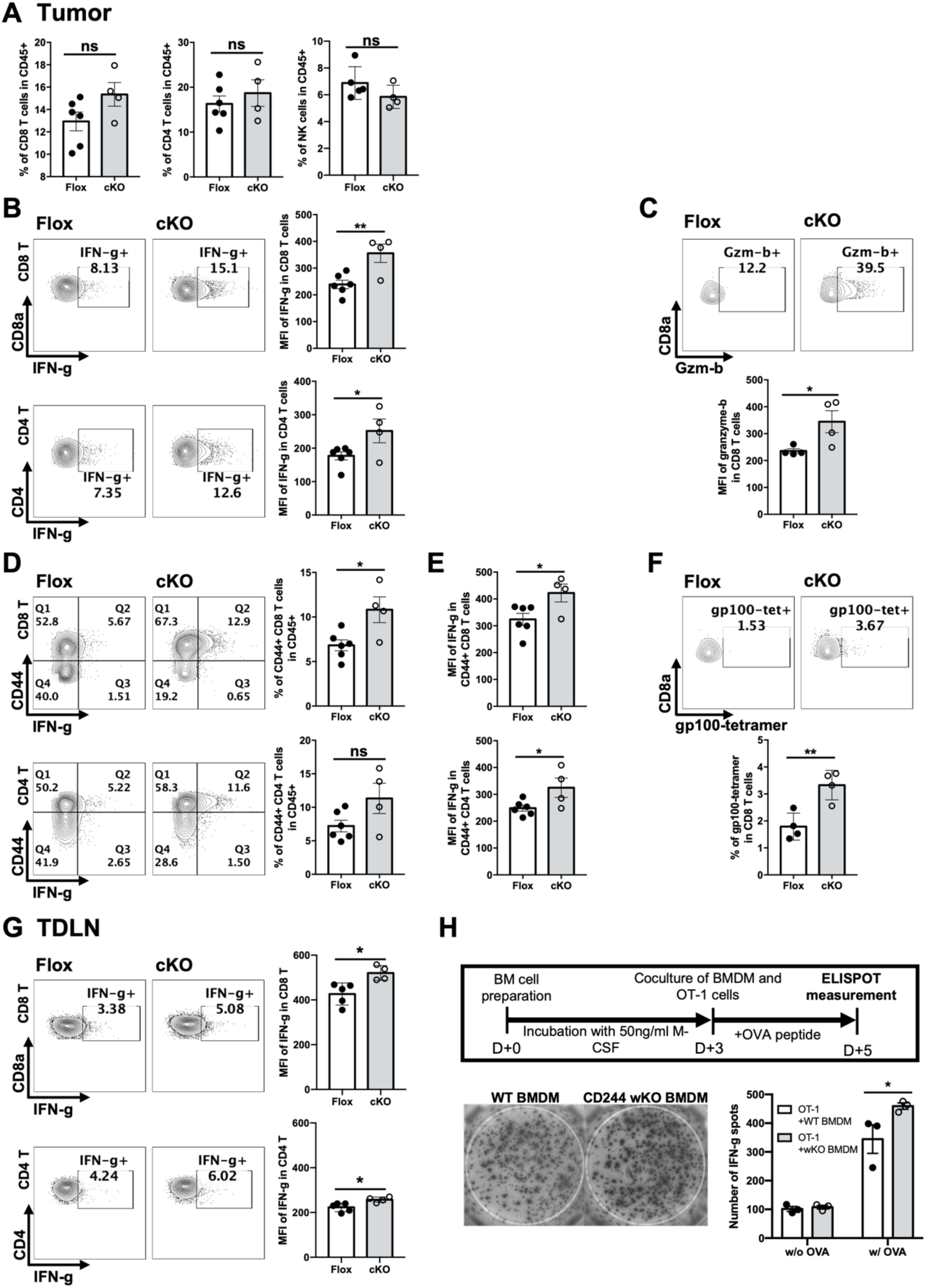
CD244 deletion on monocytes/macrophages increases antigen-specific IFN-γ secretion of T cells. (**A**) Proportion of CD8 T, CD4 T, and NK cells among total CD45+ cells in tumor mass. Unpaired Student’s *t* test; n = 6 Flox and 4 cKO (CD8 T and CD4 T cells), n = 5 Flox and 4 cKO (NK cells); N = 4. (**B–G**) IFN-γ, granzyme-b, CD44 and gp-100-specific TCR expression in tumor mass or tumor-draining lymph node (TDLN) T cells cultured with gamma-irradiated B16F10 cells. n = 4-6 Flox and 4 cKO, N = 2; unpaired Student’s *t* test. (**B**) Representative flow cytometry plot (**left**) and mean fluorescent intensity of IFN-γ (**right**) in CD8 T cells in tumor. (**C**) Representative flow cytometry plot (**upper**) and mean fluorescent intensity of granzyme-b (**lower**) in CD8 T cells in tumor. (**D**) Representative flow cytometry plot of CD44 and IFN-γ (**left**) and proportion of CD44+ CD8 T (**upper**) and CD4 T (**lower**) cells among total CD45+ cells in tumor mass (**right**). (**E**) Mean fluorescent intensity of IFN-γ in CD44+ CD8 T (**upper**) and CD4 T (**lower**) cells. (**F**) Representative flow cytometry plot (**upper**) and mean fluorescent intensity (**lower**) of H-2Db gp100-tetramer in CD8 T in CD8 T cells in tumor. **(G)** Representative flow cytometry plot (**left**) and mean fluorescent intensity (**right**) of IFN-γ expression in CD8 T **(upper)** and CD4 T **(lower)** cells in TDLN. (**G**) BMDM and OT-1 cells cocultured with cognate OVA peptide. Scheme of ELISPOT experiments **(upper)**; representative photographs of IFN-γ spots **(left)**. Number of IFN-γ spots for OT-1 cells cocultured with WT or CD244 wKO BMDMs **(right)**. n = 3, N = 2; two-way ANOVA.

To directly confirm the role of CD244-deficient macrophages in promoting antigen-specific CD8 T cell activation, we isolated CD8 T cells from the lymph nodes of OT-1 mice and incubated them with differentiated WT or CD244 wKO BMDM in the presence of ovalbumin (OVA) peptide. Consistent with our hypothesis, OT-1 cells secreted significantly more IFN-γ when cocultured with CD244 wKO BMDM than when cocultured with WT controls (Fig. 3H). Collectively, the selective deletion of CD244 in monocyte-lineage cells ameliorated tumor growth by enhancing the antigen-specific anti-tumor immunity of T cells.

### CD244 suppresses antigen presentation, phagocytosis, and autophagy in macrophages

Next, we examined whether specific deletion of CD244 in monocytes/macrophages impacts MHC class I-mediated antigen presentation. To this end, whole OVA protein was added to WT and CD244 wKO BMDM cultures for 24 h following 2 days of BM differentiation. Flow cytometry analysis with anti-H-2kb-SIINFEKL mAbs demonstrated increased surface OVA presentation in CD244 wKO macrophages compared to that of WT macrophages, without impacting the monocyte populations (Fig. 4A and B). Although the level of MHC-II surface expression was not altered (Fig. 4C), increased macrophage differentiation in CD244 wKO BMDM culture resulted in significantly more macrophages presenting OVA on MHC-I, as well as more cells expressing MHC-II molecules compared to those in the WT BMDM culture (Fig. 4D and E). Furthermore, the expression of CD80—a major costimulatory ligand that binds to the T cell costimulatory receptor CD28—was elevated in CD244-deficient macrophages within the tumor (Supplementary Fig. 2A) and BMDM (Supplementary Fig. 2B) populations. These data demonstrate that CD244 promotes tumor growth by inhibiting the differentiation and antigen presentation of monocyte lineage cells, resulting in the inhibition of antigen-specific T cell responses against tumors.

**Fig. 4.**
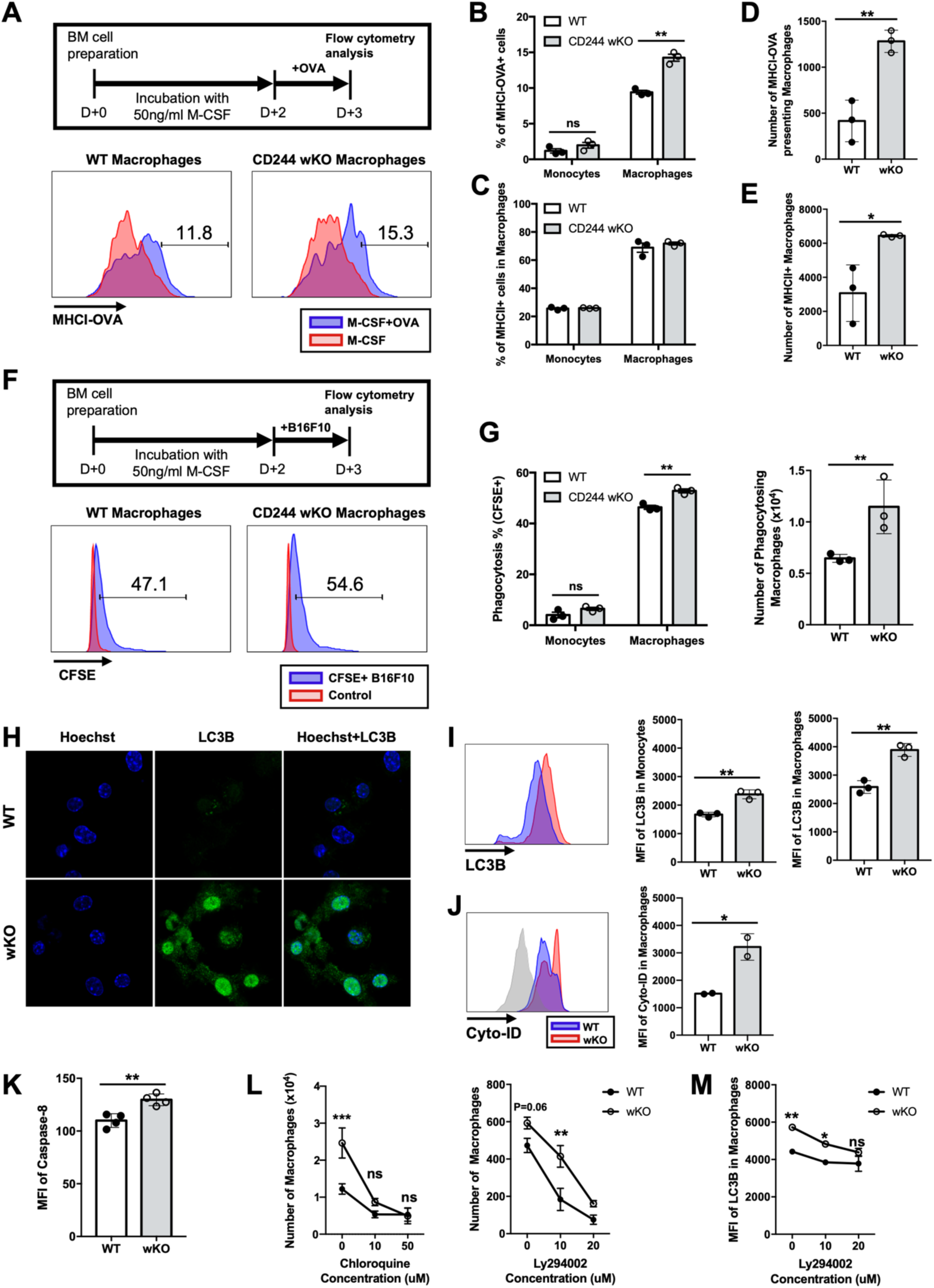
Deletion of CD244 on monocytes/macrophages increases autophagy, antigen presentation, and phagocytosis. (**A–E**) H-2kb (MHC-I)-SIINFEKL complex and MHC-II expression on D+3. n = 3, N = 3; two-way ANOVA (**B, C**) or unpaired Student’s *t* test (**D, E**). (**A**) Experiment scheme **(upper)** and representative flow cytometry plot **(lower)** of MHC-I-OVA complex expression on WT and CD244 wKO BMDM. (**B**) Proportion of MHCI-OVA+ cells in monocytes and macrophages of WT and CD244 wKO BMDM. (**C**) Proportion of MHCII+ cells in monocytes and macrophages of WT and CD244 wKO BMDM. (**D**) Absolute number of MHC-I- OVA+ macrophages in WT and CD244 wKO BMDM. (**E**) Absolute number of MHC-II+ macrophages in WT and CD244 wKO BMDM. (**F**) Experiment scheme **(upper)** and representative flow cytometry plot **(lower)** of CFSE expression on macrophages from WT and CD244 wKO BMDMs. (**G**) Proportion of CFSE+ monocytes and macrophages **(left)** and absolute CFSE+ macrophage cell number **(right)** in WT and CD244 wKO BMDMs. n = 3, N = 3. Two-way ANOVA **(left)** or unpaired Student’s *t*-test **(right)**. (**H**) Immunofluorescence of BMDMs from WT and CD244 wKO mice. (**I**) LC3B expression. Representative flow cytometry plot (**left**) and mean fluorescence intensity of LC3B in monocytes (**middle**) and macrophages (**right**). n = 3, N = 2; unpaired Student’s *t*-test. (**J**) Autophagosome and autolysosome formation. n = 2; unpaired Student’s *t*-test. (**K**) Mean fluorescence intensity of cleaved caspase-8 in D+3 BMDM. n = 3, unpaired Student’s *t*-test. (**L**) Number of macrophages in BMDM from WT and CD244 wKO mice following treatment with the autophagolysosome formation inhibitor, chloroquine **(left)** and PI3K inhibitor Ly294002 **(right)**. n = 3, N = 2; two-way ANOVA. (**M**) Mean fluorescence intensity of LC3B in D+3 WT and CD244 wKO BMDMs after addition of Ly294002. n = 3; two-way ANOVA.

Although the proinflammatory role of macrophages in the TME is highly dependent on the adaptive immune system, the importance of macrophage phagocytosis in solid and hematopoietic cancers has been highlighted ^34, 35^. To determine whether phagocytic function is also regulated by CD244, BMDMs were cocultured with γ-irradiated B16F10 cells stained with carboxyfluorescein succinimidyl ester (CFSE). The proportion of CFSE+ monocytes and macrophages were determined via flow cytometry as a measure of direct phagocytosis. CD244 wKO macrophages exhibited significantly higher phagocytic activity than WT macrophages (Fig. 4F and G). Collectively, these results indicate that CD244 signaling plays a critical role in facilitating the immunosuppressive TME by inhibiting differentiation and functional maturation of M1 macrophages in a B16F10 melanoma model.

Autophagy is a critical mechanism involved in the survival, differentiation, antigen presentation, and acquisition of phagocytic functions of monocytes and macrophages ^36–38^. Previous studies have provided evidence that CD244 as well as other SLAMF receptors contribute to the initiation step of autophagosome complex formation by binding with Beclin-1 and Vps34 ^39–41^. Hence, we propose that CD244 inhibits monocyte differentiation by regulating autophagy induction. To this end, M-CSF-treated BM cells were stained with an anti-LC3B antibody and examined by fluorescence microscopy (Fig. 4H) and flow cytometry (Fig. 4I). Results showed that CD244 wKO BMDM had significantly increased LC3 lipidation in macrophages and monocytes compared to their WT counterparts. Accordingly, CYTO-ID staining revealed that autophagosome and autophagolysosome formation was also significantly increased (Fig. 4J). Caspase activation is also important for autophagy-driven macrophage differentiation^42^. During BMDM differentiation, CD244 wKO CD11b+ cells contained significantly more cleaved caspase-8 compared with WT (Fig. 4K). To further evaluate the involvement of autophagy in the enhanced differentiation of CD244-deficeint monocytes/macrophages, we treated BMDM cultures with autophagy inhibitors. Blocking the fusion of autophagosomes and lysosomes with chloroquine in BMDM cultures significantly decreased the number of macrophages in both WT and CD244 wKO mice; however, a more significant reduction was observed in macrophages generated from CD244 wKO BM compared to that in WT (Fig. 4L, left). Treatment with the PI3K inhibitor Ly294002, which binds to Vps34 (PI3KC3), also significantly reduced macrophage differentiation in both WT and CD244 wKO mice (Fig. 4L, right). Additionally, difference between LC3B expression in WT and CD244 wKO macrophages decreased as Ly294002 concentration increased (Fig. 4M). These results demonstrate that CD244 suppresses the induction of autophagy in monocyte-lineage cells and further inhibits their differentiation into macrophages.

### CD244 deficiency in macrophages converts immunologically “cold” tumors to “hot” tumors

Our data demonstrates that CD244 suppresses anti-tumor immunity by accumulating monocytes in the TME and subsequently reducing differentiation of macrophages. In addition to their differentiation, macrophage phagocytic and antigen-presenting functions are inhibited by CD244 signaling. These data suggest that CD244-deficient macrophages could serve as a therapeutic modality to convert immunologically “cold” tumors to “hot” tumors. To assess this hypothesis, we treated monocyte-lineage specific CD244-deficient cKO mice and their control Flox mice challenged with B16 melanoma with anti-PD-L1 antibodies. Consistent with the previous finding that B16F10 tumors are largely refractory to PD-1/PD-L1 blockade therapy ^43–45^, anti-PD-L1 antibody treatment in B16F10 tumor-bearing Flox mice did not induce significant changes in tumor growth. However, anti-PD-L1 antibody treatment further decreased the size of tumors in cKO mice compared to the isotype antibody treatment (Fig. 5A and Supplementary Fig. 3A). Interestingly, memory phenotype analysis in TDLN of cKO mice revealed that the anti-PD-L1 antibody significantly increased the central memory (CD62L^+^, CD44^+^) CD8 T cell and effector memory (CD62L^-^, CD44^+^) CD4 T cell populations compared to that of anti-PD-L1 antibody-treated Flox mice and isotype-treated cKO mice (Fig. 5B and Supplementary Fig. 3B). In addition, the proportion of PD-1^+^ and T cell immunoreceptor with Ig and ITIM domains (TIGIT)^-^ CD8 T cells was significantly increased only in the tumors of cKO mice receiving the anti-PD-L1 antibody, whereas the proportion of severely exhausted PD-1 and TIGIT double-positive CD8 T cells ^4^ did not exhibit significant changes (Fig. 5C and Supplementary Fig. 3C).

**Fig. 5.**
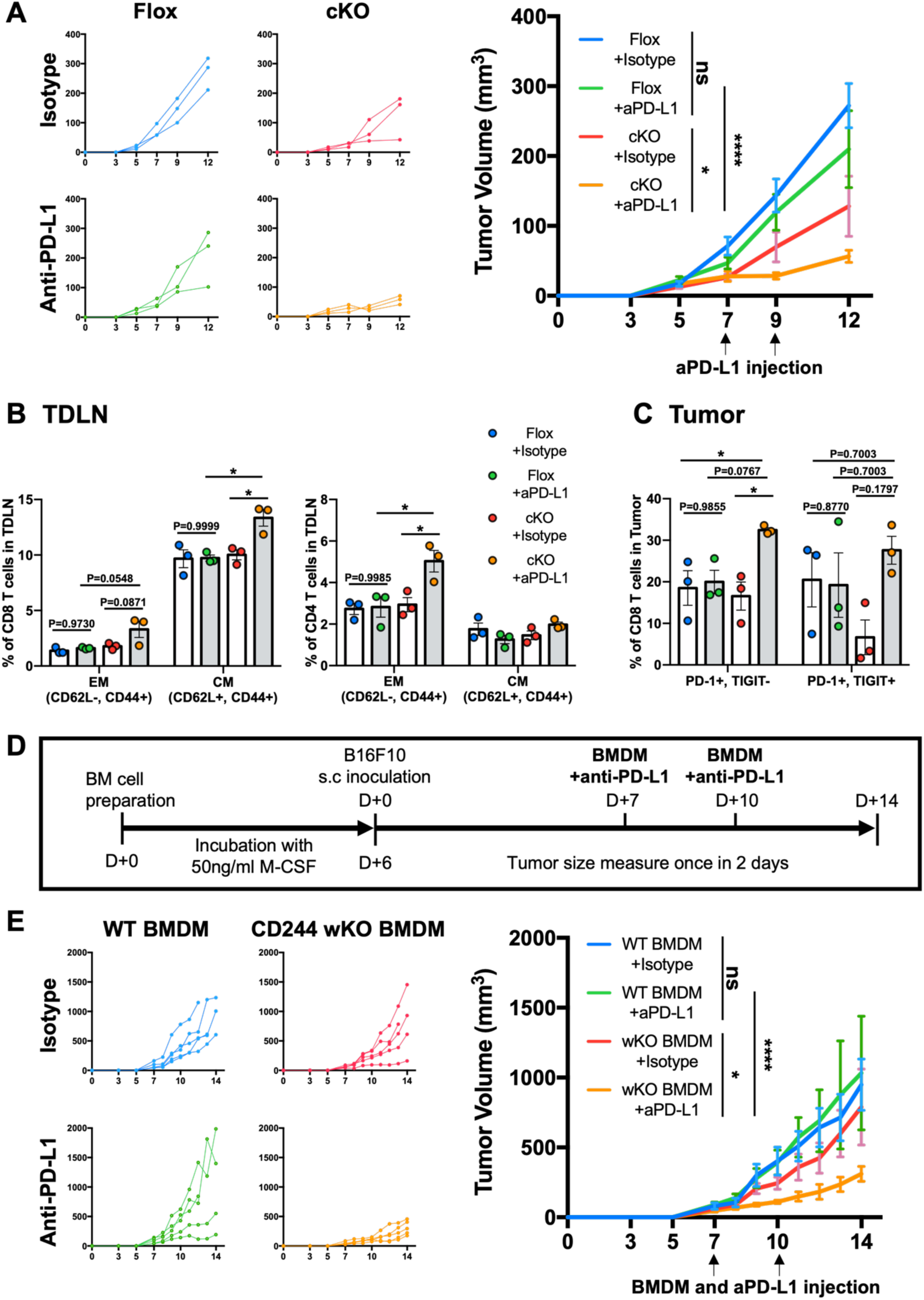
CD244-deficient macrophages significantly delay tumor growth by increasing memory T cells populations in combination with anti-PD-L1 antibody. (**A to C**) B16F10 tumor growth and intratumoral lymphocytes population were measured in anti-PD-L1 or cognate isotype antibody-treated Flox and cKO mice. n = 3, N = 2; two-way ANOVA. (**B**) CD44 and CD62L memory phenotype marker expression on CD8 T **(left)** and CD4 T **(right)** cells in the tumor draining lymph nodes (TDLN). Proportion of effector memory (CD62L^-^, CD44^+^) and central memory (CD62L^+^, CD44^+^) T cells. (**C**) Expression of PD-1 and TIGIT on tumor infiltrating CD8 T cells. Proportion of PD-1^+^, TIGIT^-^ and PD-1^+^, TIGIT^+^ cells among total CD8 T cells in tumor. (**D**) Scheme of combining BMDM and anti-PD-L1 blocking antibody therapy. (**E**) B16F10 tumor growth with WT BMDM+isotype **(upper left)**, CD244 wKO BMDM+isotype **(upper right)**, WT BMDM+anti-PD-L1 **(lower left)**, and CD244 wKO BMDM+anti-PD-L1 **(lower right)**. n = 5, N = 2; two-way ANOVA.

To assess the therapeutic potential of CD244-deficient macrophages, we performed adoptive transfer of CD244 wKO BMDM in the presence of an anti-PD-L1 antibody to B16 melanoma- bearing WT mice (Fig. 5D). While adoptive transfer of CD244 wKO BMDM into B16F10 tumor-bearing mice slightly suppressed tumor growth, the combination of CD244 wKO BMDM and anti-PD-L1 antibody significantly reduced the growth of B16F10 tumors (Fig. 5E and Supplementary Fig. 3D). These data suggest that CD244 expressed on monocytes/macrophages functions as an important checkpoint receptor within the TME, while blocking CD244 function could potently convert immunologically “cold” tumors into “hot” tumors.

### Monocytes/macrophages CD244 expression negatively correlates with patient survival

To determine the clinical significance of CD244 as an immune checkpoint receptor on monocytes/macrophages within the TME, we re-analyzed a previously reported single-cell RNA-seq dataset (SCP398 from Single Cell Portal) of CD45^+^ immune cells isolated from human melanoma tissues ^9^. Among the 16,291 cells reported in the original study, we focused on 1,391 cells annotated as monocytes and macrophages. To identify the CD244^-^ monocytes/macrophages in these cells, we further clustered them into seven subclusters (C0– 6, Fig. 6A) using the shared nearest neighbor clustering method in Seurat. CD244 expression was categorized into the following three groups (Fig. 6B):1) C0–1 with relatively high enrichment of CD244 expression, 2) C2–3 with low enrichment of CD244 expression, and 3) C4–6 with no expression of CD244. Of these groups, we focused on C0–1, with significant numbers of both CD244^+^ and CD244^-^ monocytes/macrophages, for a fair comparison between the two cell types. A total of 511 and 221 genes were identified as predominantly upregulated in CD244^+^ and CD244^-^ monocytes/macrophages, respectively (Fig. 6C). The 221 genes predominantly expressed in CD244^-^ monocytes/macrophages were primarily associated with phagocytosis (phagosome), antigen processing and presentation, and autophagy (Fig. 6D and E), which is consistent with the results found in cKO mice challenged with B16F10 melanoma.

**Fig. 6.**
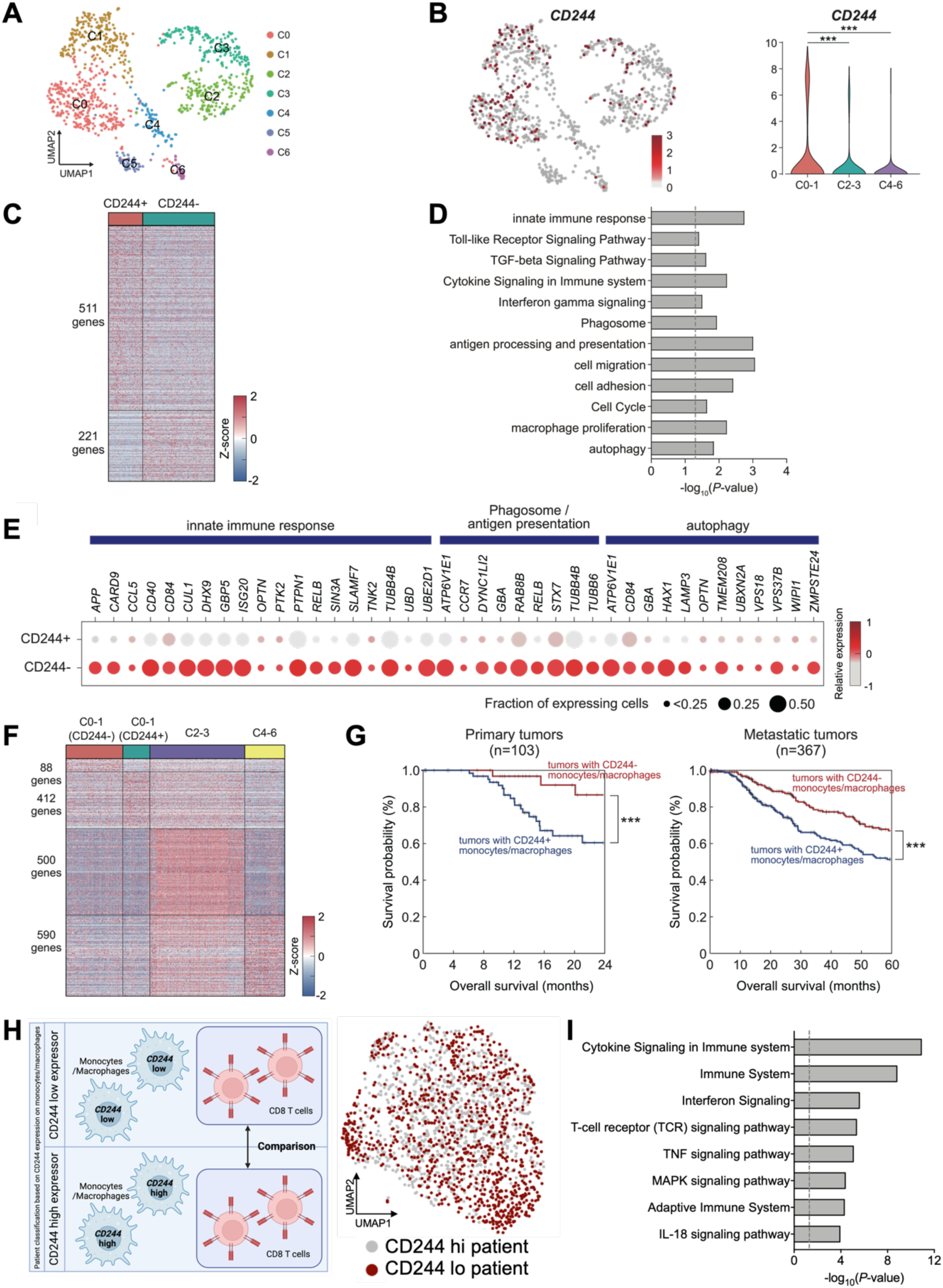
scRNA seq data and deconvolution analysis reveals that CD244 regulates the fate of monocytes/macrophages and melanoma patients’ survival. (**A**) Uniform manifold approximation and projection (UMAP) plot showing seven subclusters (C0–6) of monocytes/macrophages identified from single cell RNA-seq data. (**B**) UMAP showing CD244^-^expressing monocytes/macrophages (red highlighted cells). (**right**) Distribution of CD244 expression levels in three cell groups (C0–1, C2–3 and C4–6). \*\*\**P* < 1.010^-3^ from non-parametric one-way ANOVA with Tukey’s post-hoc correction. (**C**) Differentially expressed genes between CD244^+^ and CD244^-^ monocytes/macrophages in C0–1. Color bar, gradient of Z-scores with zero mean and unit standard deviation across cells in C0–1 for individual genes. (**D**) Cellular pathways enriched by the 221 genes upregulated in CD244^-^ monocytes/macrophages; presented as –log_10_ (*P*-value). (**E**) Preferential expression of genes involved in the innate immune response, phagosome/antigen presentation, and autophagy in CD244^-^ monocytes/macrophages. Color bar, gradient of gene Z-scores. (**F**) Genes predominantly upregulated in CD244^+^ and CD244^-^ monocytes/macrophages in C0–1, C2–3 and C4–6. Color bar, Z-score gradient. (**G**) Overall survival of patients with primary (**left**) or metastatic (**right**) tumors, with and without CD244^-^ monocytes/macrophages. (**H**) Schematic representation of patients’ classification based on CD244 expression level of G3 monocytes/macrophages (**left**). UMAP plot showing CD8 T cells of CD244 low and high patients (based on monocytes/macrophages; **right**) (**I**) Cellular pathways enriched by the genes upregulated in CD8 T cells of CD244 low patients; presented as –log_10_ (*P*-value).

The presence of CD244^-^ monocytes/macrophages in human melanoma tissue prompted us to assess the clinical implications of CD244 on monocytes/macrophages in the pathogenesis of human melanoma. To this end, we obtained bulk RNA-seq datasets generated from the tumor tissues of 470 melanoma patients (103 primary and 367 metastatic tumors)^46^ from The Cancer Genome Atlas (TCGA) database (TCGA-SKCM). We then estimated the proportions of the following four cell types identified from the two bulk RNA-seq dataset analyses: 1 & 2) CD244- and CD244+ cells in C0–1, 3) cells in C2–3, and 4) cells in C4–6. We then identified genes that were predominantly upregulated in the four cell types (Fig. 6F). Notably, 511 and 221 genes identified from C0–1 only (Fig. 6C) were reduced to 412 and 88 genes, respectively, when we further filtered out the genes exhibiting increased expression in cells in C2–3 or C4– 6 to ensure the specificity of their expression in C0–1 compared to C2–6. The proportions of the four cell types in individual tumor tissues from TCGA dataset were estimated using CIBERSORTx ^47^, with the genes upregulated in each cell type. We then compared the survival of patients with and without CD244^-^ monocytes/macrophages and found that patients with CD244^-^ monocytes/macrophages showed better overall survival than those without CD244^-^ monocytes/macrophages in two different patient cohorts with primary (2-year survival; Fig. 6G, left) and metastatic (5-year survival; Fig. 6G, right) tumors. These data are consistent with our results showing delayed tumor growth in monocytes/macrophages-specific CD244- deficient mice (Fig. 1F and Supplementary Fig. 5B and C).

Then we wondered CD244 low monocytes/macrophages can enhance antigen-specific T cell activity in human same as in mice model. We classified patients into CD244 high and CD244 low groups according to their average expression level of CD244 on G3 monocytes/macrophages and compared their CD8 T cells, regardless of CD244 expression in T cells. (Fig. 6H and Supplementary Fig. 5A). We obtained 207 genes predominantly upregulated in CD8 T cells from CD244 low patients and these DEGs were primarily associated with cytokine signaling, TCR signaling and adaptive immune system (Fig. 6I). Together, these data suggest that CD244 functions as a novel immune checkpoint receptor on monocytes/macrophages in patients with melanoma.

## DISCUSSION

Although studies have underscored the role of CD244 expressed on lymphoid cells in cancer, its precise role in myeloid cells has remained elusive, as most studies have been confined to evaluating the correlation of its expression and function in specific disease settings. By taking several measures, we present direct evidence that CD244 expressed on monocyte-lineage cells serves as an immune checkpoint receptor within the TME. More specifically, our LysM-cre^+/-^ CD244^fl/fl^ cKO mice challenged with B16 melanoma showed delayed tumor growth compared to WT controls, through enhanced anti-tumor T cell immunity. Importantly, CD244 cKO mice exhibited synergistic reduction with anti-PD-L1 antibody treatment of B16 melanoma, which was largely refractory to monotherapy ^43–45^. Furthermore, adoptive transfer of CD244-deficient BMDM into B16F10 tumor-bearing WT mice resulted in slight retardation of tumor growth; however, the combination of CD244-deficient BMDMs and anti-PD-L1 antibody significantly reduced the growth of B16F10 melanoma by restoring and preventing exhaustion of T cell immunity. At the molecular level, CD244 signaling suppresses autophagy and autophagosome formation, leading to the inhibition of monocyte-to-macrophage differentiation, antigen presentation, and tumor phagocytosis of CD11b^+^Ly6C^low^F4/80^+^ macrophages. Consistent with the results in the mouse model, our cell-type deconvolution analysis highlights the potential implications of CD244^-^ monocytes/macrophages in the survival of patients with melanoma, thereby providing significant insights into the development of novel therapeutic modalities targeting CD244 within the TME.

SLAM family receptors (SFR) include six transmembrane receptors: SLAM (SLAMF1; CD150), CD244 (SLAMF4; 2B4), Ly-9 (SLAMF3; CD229), CD84 (SLAMF5), SLAMF6 (Ly108; NTB-A), and SLAMF7 (CRACC; CS1) ^48–50^. All SFRs, except CD244, are homotypic receptors that engage in either trans-interactions on neighboring cells or cis interactions on the same hematopoietic cells. CD244 interacts with CD48 (SLAMF2), the expression of which is restricted to hematopoietic cells. Although extensive studies by Veillette et group ^51–54^ and our group ^13, 55–59^ have revealed the importance of CD244 as a self-tolerance receptor on NK cells, the role of SLAM family receptors on monocytes/macrophages has only recently been recognized ^27, 29, 40, 60, 61^. Chen et al. demonstrated that SLAMF7 promotes phagocytosis of hematopoietic tumor cells in macrophages in the absence of SIRPa-CD47 interactions ^35^. Moreover, Li et al. unexpectedly found that CD244 negatively regulates LRP1-dependent, CD47-independent phagocytic pathways in macrophages against hematopoietic cells ^61^. Their conclusion was based on an increased ability of CD244+ WT macrophages to clear autologous CD48 KO hematopoietic tumor target cells. However, interestingly, SFR wKO mice did not reject non-hematopoietic skin grafts, suggesting that CD244 likely restricts macrophage phagocytosis against hematopoietic tumors, but not in solid tissue or organs. Our data agree with their results in that CD244 serves as an important self-tolerance receptor that inhibits phagocytosis pathways in macrophages but further extends to solid cancer by generating a cold TME via regulation of phagocytic function acquisition.

In addition to its role in regulating phagocytosis, we demonstrate that CD244 inhibits class I-dependent antigen presentation in CD11b^+^Ly6C^low^F4/80^+^ macrophages, leading to insufficient T cell activation. Besides CD103+ DCs present inside the tumor that can properly activate CD8 T cells through cross-presentation ^62, 63^, this ability has also been reported for M1 macrophages^64^. Consistent with these findings, our data highlight the role of CD244-deficient M1 macrophages in activating tumor-specific CD44+ memory CD4 and CD8 T cells and triggering IFN-γ secretion in an antigen-dependent manner. Therefore, CD244 serves as an important immune checkpoint receptor by regulating both the innate and adaptive immune axes in monocytes/macrophages within the TME. Our data also provide a mechanistic basis underlying the immunosuppressive role in both mouse and human head and neck squamous cell carcinoma models ^29^. In these tumor models, CD244 expression in head and neck squamous carcinoma is positively correlated with PD-1 and PD-L1 expression, as well as that of immunosuppressive genes including *ARG, IL10*, and *TGFΒ1* ^29^. The upregulation of inhibitory genes in CD244^+^ myeloid cells might also have contributed to the therapeutic synergy of CD244-deficient BMDM and anti-PD-L1 antibody, as shown in our studies with the B16 melanoma model. Interestingly, CD244 expression has also been associated with the expression of other checkpoint receptors, including CTLA-4, TIM-3, and LAG-3 on CD8+ T cells ^31^, highlighting the multi-level regulation of tumor immunity. Therefore, the synergistic blockade of pathways preventing the upregulation of these co-inhibitory receptors would prove beneficial in preventing immune exhaustion within the TME.

Given that CD244 on monocytes/macrophages regulates both the innate and adaptive axes of tumor immunity, elimination of CD244 on BMDM could lead to the conversion of immunologically cold microenvironments into more proinflammatory environments (hot tumors). Flow cytometric analysis in cKO mice showed that anti-PD-L1 antibody increased the memory phenotype of CD8/CD4 T cells and PD-1^+^ TIGIT^-^ CD8 T cells without altering the proportion of PD-1/TIGIT double-positive CD8 T cells. TIGIT expression is largely dependent on the exhaustion status of CD8 T cells, which is inversely correlated with effector function in various tumors, including melanoma ^65–68^. In addition, TIGIT^+^ T cells co-express numerous other exhaustion markers, including TIM-3 and LAG-3, thus representing severely exhausted T-cells ^4, 69^. These data demonstrate that removal of CD244-mediated inhibitory signals on macrophages alone is not sufficient to trigger strong tumor immunity in immunologically cold B16 melanoma *in vivo*; however, concomitant blocking of PD-1/PD-L1 interactions on T cells releases the break on exhausted T cells and restores anti-tumor effector functions. Therefore, our results present CD244-deficient BMDM with checkpoint blockade as a promising therapeutic combination modality in patients with immunologically cold tumors. In this context, simultaneous blockade of the SIRPa-CD47 “don’t-eat-me signals” interaction and CD244/CD48 could lead to synergistic activation of macrophage-based tumor immunity ^70^, while also enhancing effector functions of exhausted CD8 T cells in patients with cancer. Taken together, our data shed light on the development of novel combinatorial therapies against relapsed and resistant immunologically cold tumors.

Finally, the cell-type deconvolution analysis suggested the potential implications of CD244^-^ monocytes/macrophages in the survival of patients with melanoma. The increased proportion of CD244^-^ monocytes/macrophages was associated with improved survival in patients with both primary and metastatic tumors, suggesting that the presence of CD244^-^ monocytes/macrophages could be a prognostic factor in these patients. Additionally, analysis of scRNA-seq data from melanoma patients revealed that autophagy, antigen presentation, and phagocytosis pathways are upregulated in CD244- monocytes/macrophages, implying that CD244 initiates tumor immune escape in patients with melanoma by regulating multiple autophagy-mediated functional axes.

Certain limitations were noted in this study that warrant further investigation. First, although the primary target of LysM-cre is monocytes/macrophages, it is known to also target neutrophils with high efficiency and DCs with low efficiency. However, we did not observe significant changes in the CD244 levels or proportion within these cell populations. In the case of neutrophils, there was no significant increase in CD244 expression in tumors compared to that in the spleen; hence, we postulate that CD244 on neutrophils does not have a tumor-specific role. Second, given that M1 macrophages are commonly involved in the development and progression of all tumor types, it is also necessary to investigate whether the anti-tumorigenic function of CD244^-^ monocyte lineage cells is effective in other tumors. These investigations may provide an important therapeutic modality that can be applied for the treatment of a broad spectrum of human cancers.

In summary, our results in mouse melanoma models as well as human patients underscore a previously unidentified role of CD244 on monocytes/macrophages as a novel immune checkpoint receptor, potentially contributing to T cell exhaustion and immunologically ‘cold’ microenvironments. Therefore, CD244-deficient macrophages could serve as a therapeutic modality to convert immunologically “cold” tumors to “hot” tumors, which when combined with checkpoint blockades can synergistically increase anti-tumor efficacy. Furthermore, the proportions of CD244^+^ and CD244^-^ monocyte lineage cells can serve as negative or positive prognostic markers in patients, thereby tailoring new potential therapeutic options for melanoma.

## MATERIALS AND METHODS

### Study design

This study was a controlled laboratory experiment using a mouse model and primarily aimed to identify the role of CD244 expressed on monocytes/macrophages in the TME. The sample size applied for the mouse tumor implantation experiments was determined to achieve a significance level greater than 95% using G-power software. No outliers were excluded from the experiments or analyses reported in this study. The final endpoint of the mouse tumor growth experiment was determined according to the guidelines of the Institutional Animal Care and Use Committee of the Korea University. For all the studies, biological replicates and the number of independently performed experiments are indicated in the figure legends. The sum of the data units represents the number of biological replicates, which are depicted as individual values in the bar graphs. The experiments were not randomized. The investigators were not blinded to allocation during the experiments or outcome assessment.

### Mice

All animal experiments were approved by the Institutional Animal Care and Use Committee of Korea University (approval number: KUIACUC-2019-0101) and followed the guidelines and regulations of the Institutional Animal Care and Use Committee of Korea University. WT C57BL/6 mice were purchased from Orient Bio. Inc. (Seongnam, Korea). CD244^-/-^ mice on a C57BL/6 background were generated as previously described ^13^. CD244^f/f^ mice were generated using Cyagen Biosciences (Santa Clara, CA, USA). CD244^f/f^ mice were bred with LysM^cre+^ (B6.129P2-LysMtm1(cre)Ifo/J) mice from Jackson Laboratory (Bar Harbor, ME, USA) to generate CD244^fl/fl^ and LysM-cre^+/-^CD244^fl/fl^ mice, and littermates were used for conditional CD244-deficient mice experiments. Female mice between 5 and 10 weeks of age were used in the experiments. Mice were bred and maintained in a specific pathogen-free facility at Korea University.

### Cell lines and tumor models

B16F10 melanoma cells were purchased from American Type Culture Collection (ATCC). To establish subcutaneous tumors, 1 × 10^6^ B16F10 cells were injected into right flank of mice, which formed a tumor with a 1-cm diameter within 1–3 weeks of injection. Tumors were measured regularly with digital callipers and tumor volumes were calculated by the fomula: length×width×height/2. Anti-PD-L1 andithody (200ug, clone 10F.9G2, Bio X Cell, West Lebanon, NH, USA) or rat IgG2b isotype control antibody (200ug, clone LTF-2, Bio X Cell, West Lebanon, NH, USA) was injected intraperitoneally 2 times in 100ul PBS on days 7, 10 after inoculation of tumor cells.

### Cell isolation

Single cell suspensions were prepared from the spleen and BM, followed by red blood cell removal using ammonium chloride lysis buffer. Monocytes were purified from the BM using the Monocyte Isolation Kit (Miltenyi Biotec, Auburn, CA, USA) according to the manufacturer’s instructions. Single cell suspensions from tumor tissues were prepared using the Mouse Tumor Dissociation Kit (Miltenyi Biotec, Auburn, CA, USA) and Percoll density gradient separation according to the manufacturer’s recommendations (GE Healthcare, Chicago, IL, USA).

### Isolation of CD11b+ cells and monocytes

Single-cell suspensions were prepared from the tumor and BM as described above. For CD11b+ cell isolation, enriched single-cell suspensions from tumor lysates were labelled with biotinylated anti-CD11b antibody (BioLegend, San Diego, CA, USA) and separated on MACS columns (Miltenyi Biotec, Auburn, CA, USA). The monocytes were sorted using a monocyte isolation kit (Miltenyi Biotec, Auburn, CA, USA).

### BMDM and tumor-infiltrating monocytes differentiation

Enriched monocytes, whole BM cells, or CD11b+ enriched single cell suspensions of B16F10 tumors were cultured in the presence of 50 ng/mL recombinant M-CSF (Peprotech, Cranbury, NJ, USA) in RPMI (Wellgene, Gyeongsan-si, Korea) supplemented with 10% FBS, 10 mM HEPES, 20 µM 2-mercapoethantol, 1% penicillin/streptomycin, and 1% non-essential amino acids.

### Flow cytometry

Typically, up to 1 × 10^6^ cells were incubated with Fc-block for 5 min at room temperature (RT) and surface staining was performed for 30 min at 4 °C in the dark, followed by centrifugation at 1700 rpm for 5 min. H-2Db gp100 Tetramer-EGSRNQDWL-PE (MBL International, Woburn, Massachusetts, USA) staining was performed for 30 min at 4 °C in the dark, prior to surface staining. For intracellular staining, CD45+ enriched single cell suspensions of tumor or tumor draining lymph nodes were incubated for 16 h with γ-irradiated B16F10 melanoma and brefeldin A. After incubation, cells were surface stained and then perforated and intracellularly stained with a BD fixation/permeabilization kit (BD Biosciences, San Jose, CA, USA) according to the manufacturer’s recommendations. Cells were run on a FACSCanto II flow cytometer (BD Biosciences, San Jose, CA, USA), and data were analyzed using FlowJo software (BD Biosciences, San Jose, CA, USA). A list of the antibodies used is provided in Supplementary Table 1.

### Quantitative PCR

Total RNA was extracted using TRIzol reagent (Invitrogen, Middlesex County, MA). cDNA was synthesized using the TOPscript^TM^ cDNA Synthesis Kit (Enzynomics, Daejeon, Korea). Real-time PCR was performed using SYBR Green (Bio-Rad) on a StepOnePlus^TM^ (Applied Biosystems, Middlesex County, MA). Gene expression was normalized to the level of GAPDH mRNA, and relative expression levels were calculated according to the 2^-ΔΔCt^ method. Genes were amplified using the primers listed in Supplementary Table 2.

### ELISPOT

After 3 days of BMDM differentiation, BMDMs and 5 × 10^4^ of lymph node cells from OT-1 transgenic mice were cocultured at various ratios with cognate OVA peptide (SIINFEKL; 1 µg/mL). Cells were plated in the Mouse IFN-γ ELISPOT^PLUS^ kit (ALP) (MABTECH, Nacka Strand, Sweden) and IFN-γ spots were detected after 48 h using ELISPOT reader systems (Autoimmun Diagnostika GmbH, Strassberg, Germany).

### Antigen presentation and phagocytosis assay

After 2 days of BMDM differentiation, 1 mg/mL of whole OVA protein (Invivogen, San Diego, CA, USA) or γ-irradiated and CFSE stained B16F10 melanoma cells were added to the culture. OVA peptide-conjugated MHC-I complex and MHC-II were labeled with anti-H2-kb-SIINFEKL antibody and anti-MHC II antibody (BioLegend, San Diego, CA, USA), respectively to assess antigen presentation and CFSE fluorescence was measured to assess phagocytosis; expression was measured using a FACSCanto II flow cytometer (BD Biosciences, San Jose, CA, USA).

### Fluorescence microscopy

BMDM cultured on confocal dishes were fixed and permeabilized with 4.2% paraformaldehyde solution and stained with anti-LC3B antibody (1:200) and Hoechst 33342 (1:1000). After 2 h of incubation, the cells were washed and stained with Alexa555-rabbit IgG secondary antibodies and incubated for 1 h at RT. All images were captured using an LSM700 confocal laser-scanning microscope (Carl Zeiss, Oberkochen, Germany) equipped with a 63× oil-immersive lens.

### scRNA-seq data analysis

A scRNA-seq dataset (accession ID: SCP398) for patients with melanoma that was previously reported ^9^ was obtained from the Single Cell Portal (https://singlecell.broadinstitute.org/single_cell). From the dataset, we collected unique molecular identifier (UMI) count matrix for cells selected based on the quality control criteria used in the original study ^9^ as well as the cell type annotations reported in the original study. Only UMI count profiles for 1,391 cells corresponding to monocytes and macrophages, among the 16,291 cells in the dataset, were used for our analysis. The count matrix was normalized by cell-specific size factors using the Seurat (v4.0.6) R package ^71^ and subsequently log_2_- transformed after addition of a pseudo-count of 1. Highly variable genes (HVGs) were identified using FindVariableFeatures in Seurat with an FDR < 0.05. Using these HVGs, we clustered 1,391 cells corresponding to monocytes and macrophages with the first 50 principal components (PCs) obtained for the HVGs using FindClusters function in Seurat. We then visualized the resulting subclusters of monocytes and macrophages using uniform manifold approximation and projection (UMAP) with the RunUMAP function in Seurat. Genes predominantly upregulated in each subcluster were identified using the Wilcoxon rank sum test with the FindAllMarkers function in Seurat and an adjusted *P*-value < 0.05, and log_2_-fold change > 0.25. In addition, differentially expressed genes (DEGs) between CD244^+^ and CD244^-^ cells were identified using the same function. Functional enrichment analysis of DEGs was performed using ConsensusPathDB software (version 35) ^72^. Gene ontology biological processes (GOBPs) and pathways enriched by the genes were identified as those with *P* < 0.05.

### Bulk RNA-seq data analysis

We obtained two RNA-seq datasets of melanoma patients previously reported ^46, 73^ from TCGA database (TCGA-SKCM; https://portal.gdc.cancer.gov) and the GEO database (accession ID: GSE91061). We first estimated the proportions of CD244^+^ and CD244^-^ cells in individual melanoma tissue samples through cell deconvolution analysis using CIBERSORTx software with absolute mode and disabled quantile normalization ^47^. The fragments per kilobase of transcript per million mapped reads (FPKM) values of individual melanoma tissue samples were uploaded to CIBERSORTx as a mixture file. For survival analysis, the patients were divided into two groups: with and without CD244^-^ cell fractions. The cumulative event (death) rate was calculated for each patient group using the Kaplan–Meier method, and the survival curves of the two patient groups were compared using the Kaplan–Meier (log rank) test.

### Statistical analyses

All data are presented as mean ± standard error of the mean (S.E.M). Details on the sample size (biological replicates), number of repetitions, and statistical tests are listed in the figure legends. Student’s *t*-test or Mann-Whitney *U*-test was used to determine statistical significance between the two groups. Two-way ANOVA with correction for multiple comparison tests was used to determine significant differences between groups when more than one variable was being assessed. Significance was defined at *P* < 0.05. All analyses were performed using the Prism software (version 7.0; GraphPad Software, Inc.).

## Author contributions

Conceptualization: KML, TJK, JSK

Methodology: JSK, TJK, SHC, KML, DHH, JWK, KTI, MHS, IMR, SIP

Investigation: JSK, TJK, SHC, SHP, HJH, YJP, JMK, JYK, CJY, EJL, HML, DHK, KHL

Visualization: JSK, SHC Funding acquisition: KML

Project administration: KML, TJK, JSK

Supervision: KML, TJK, DHH, KHL

Writing – original draft: JSK, KML, SHC

Writing – review & editing: JSK, KML, TJK, DHH, KHL, JWL, IMR, SIP

## Data and materials availability

All data associated with this study are presented in the paper or in the Supplementary Materials. All resources generated in this study are available upon request from the lead author upon providing a complete material transfer agreement. The accession number for the data that we demonstrated here is an scRNA-seq <u>dataset</u> (accession ID: SCP398), RNA-seq datasets of melanoma patients previously reported from TCGA database (TCGA-SKCM; https://portal.gdc.cancer.gov), and GEO database (accession ID: GSE91061). All source codes have been deposited at https://doi.org/10.5281/zenodo.6844694 and are publicly available at the date of publication.

---

**Fig. S1.**
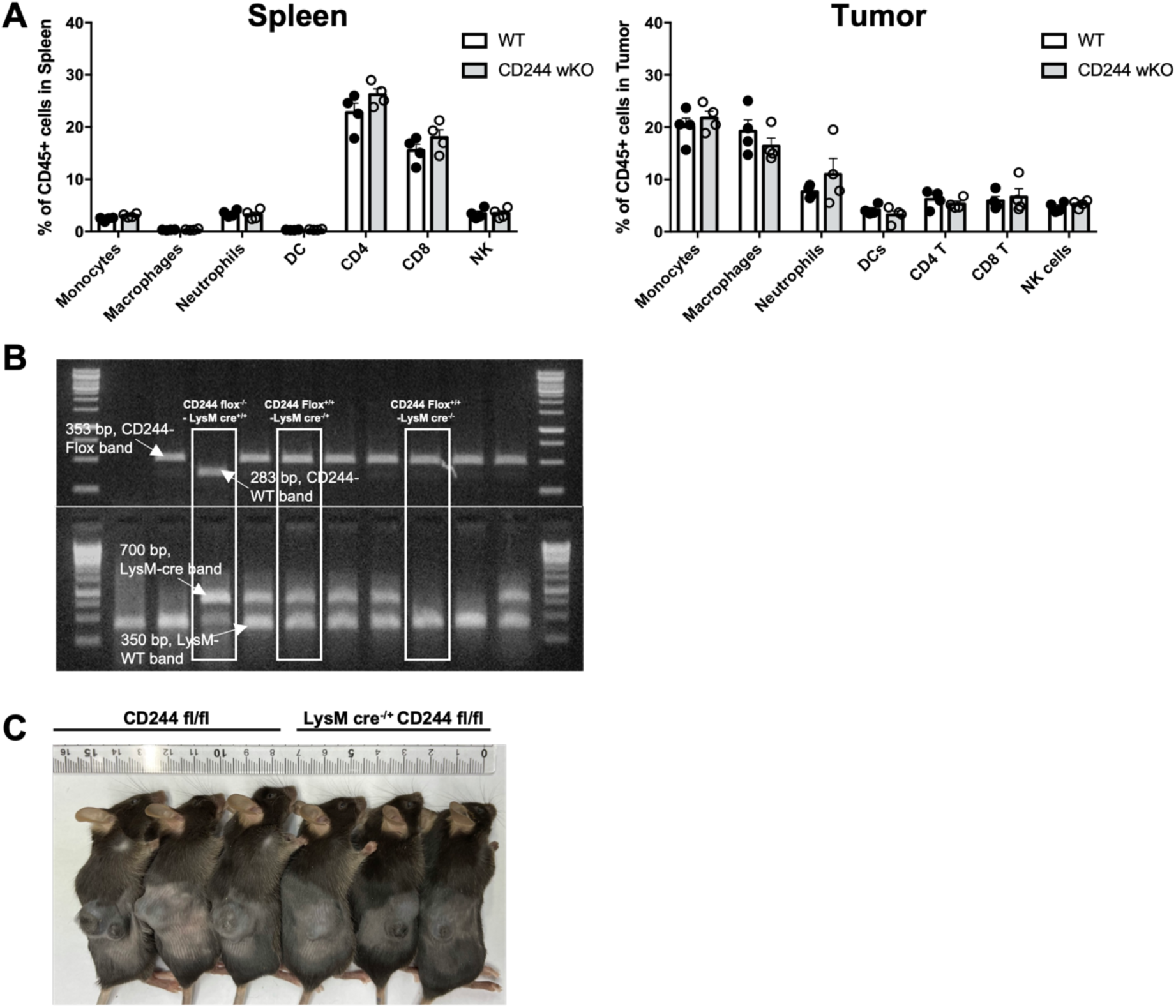
Proportion of immune cell subtypes in spleen and tumor of tumor bearing CD244 whole KO mice and generation of conditional KO mice. (A) Proportion of immune cell subtypes in spleen and tumor of tumor bearing WT and CD244 whole KO mice. (B) Genotyping results of CD244^fl/fl^ (Flox) and LysMcre^-/+^ CD244^fl/fl^ (cKO) mice. (C) Representative photograph of B16F10 inoculated Flox and cKO mice at D+14.

**Fig. S2.**
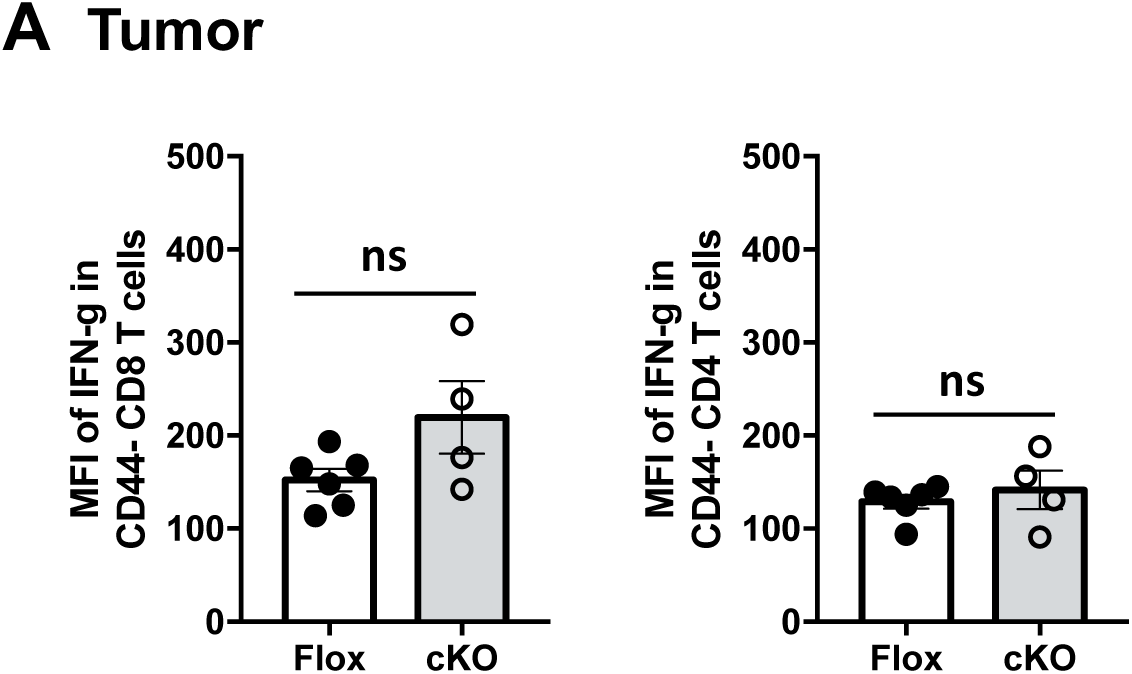
IFN-γ expression in tumor infiltrating CD4 T cells from Flox and cKO mice. (A) Mean fluorescent intensity of IFN-γ (**right**) in CD44- CD8 T and CD4 T cells in tumor.

**Fig. S3.**
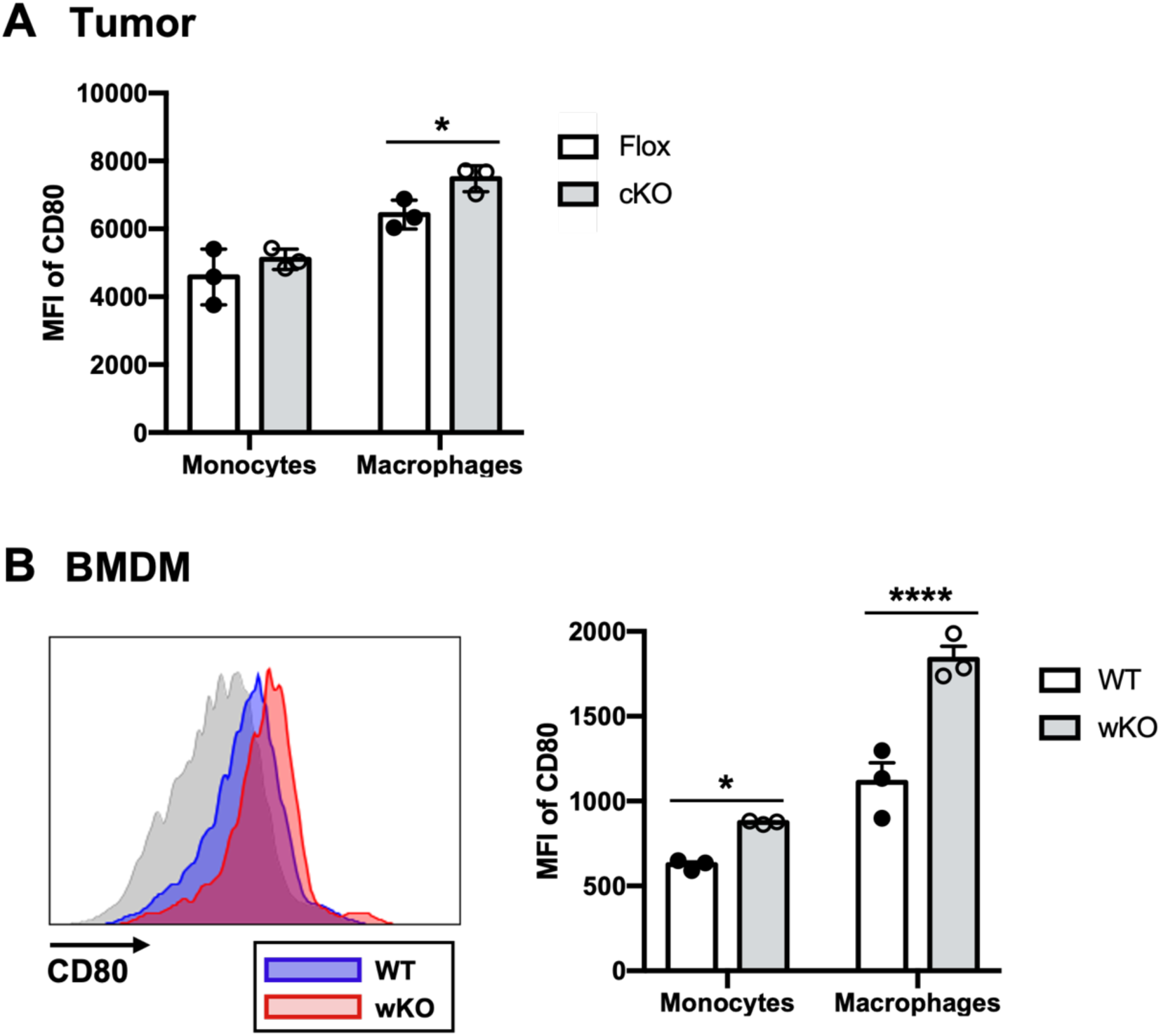
CD80 co-stimulatory molecule expression on monocytes/macrophages. (A) Mean fluorescent intensity of CD80 on tumor infiltrating monocytes and macrophages. n=3, N=2; two-way ANOVA. (B) Representative flow cytometry histogram (left) and mean fluorescent intensity of CD80 on monocytes and macrophages from D+3 BMDM culture. n=3, N=2; two-way ANOVA.

**Fig. S4.**
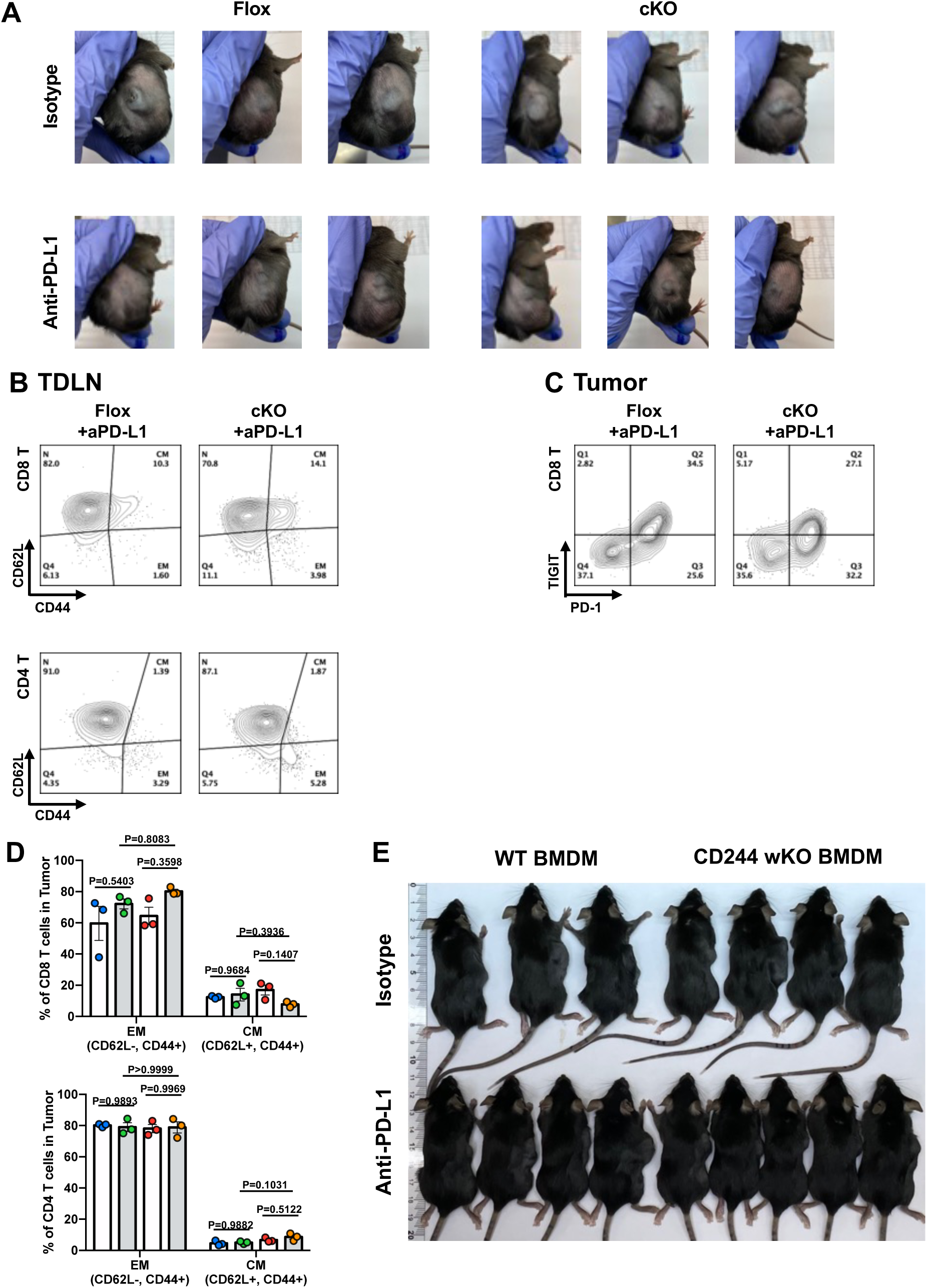
Anti-PD-L1 antibody augmented anti-tumor activity of CD244-deficient macrophages. (A) Representative photograph of B16F10 inoculated Flox and cKO mice treated with anti-PD-L1 or cognate isotype antibody. n = 3, N = 2. (B) Representative flow cytometry plots of CD44 and CD62L on CD8 T cells (**upper**) and CD4 T cells (**lower**) in tumor-draining lymph nodes. (C) Representative flow cytometry plots of PD-1 and TIGIT on CD8 T cells in tumor. (D) CD44 and CD62L memory phenotype marker expression on CD8 T **(upper)** and CD4 T **(lower)** cells in the tumor. Proportion of effector memory (CD62L^-^, CD44^+^) and central memory (CD62L^+^, CD44^+^) T cells. n = 3, N = 2; two-way ANOVA. (E) Representative photograph of B16F10 inoculated Flox and cKO mice treated with WT or CD244 wKO BMDM and anti-PD-L1 or cognate isotype antibody. n = 3-5, N = 2.

**Fig. S5.**
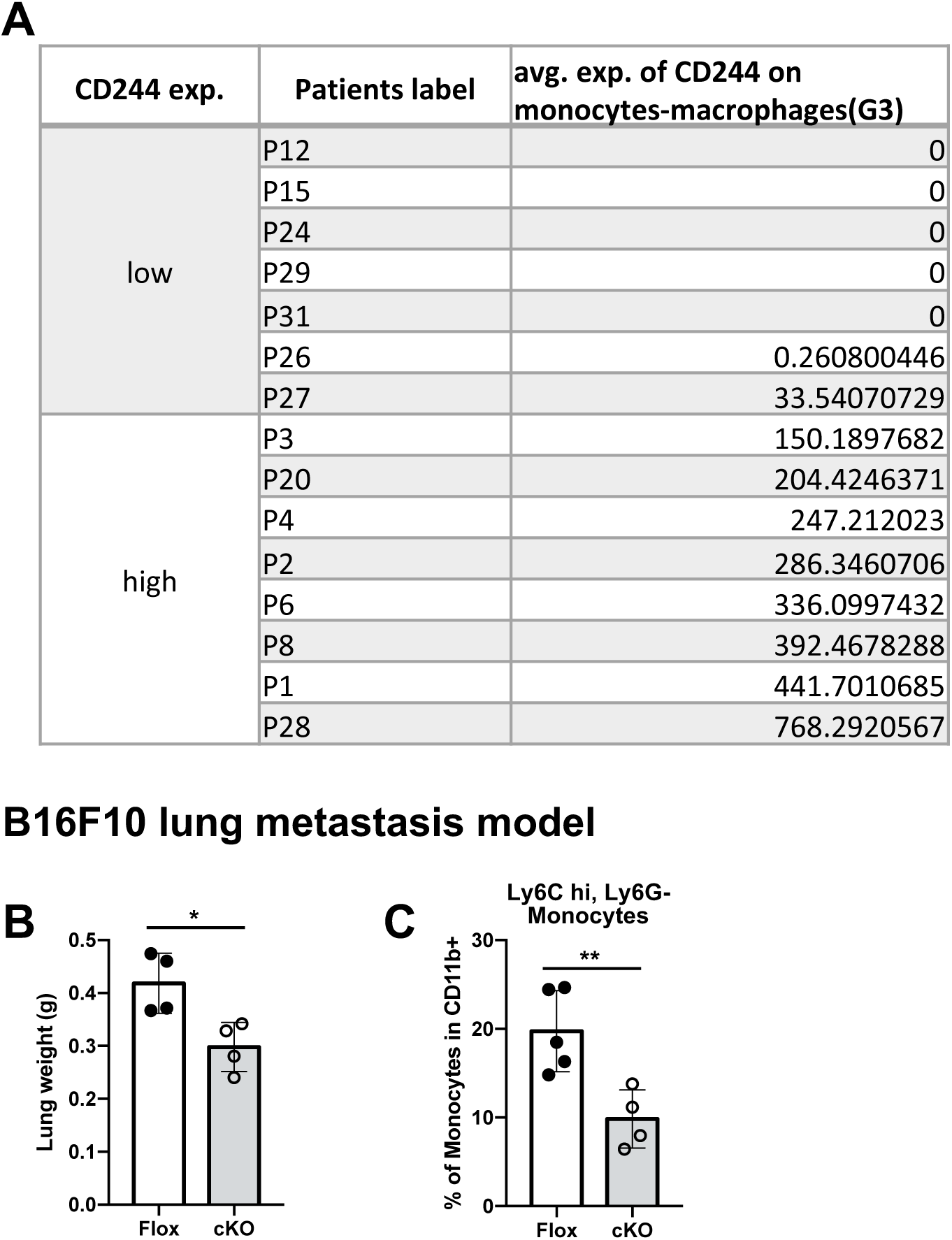
Absence of CD244 decreased tumor burden in B16F10 lung metastasis model and decreased monocytes accumulation same as subcutaneous model. (A) Average expression of CD244 on monocytes-macrophages cluster (G3) in each patient. (B) Lung weight measured at 14 days after intravenous injection of B16F10 into Flox and cKO mice. (C) Proportion of CD11b^+^, Ly6G^-^, Ly6C^hi^, F4/80^lo^ monocytes in lung.

**Table. S1.** Flow cytometry antibody list

| **Flow cytometry antibodies** |  |  |  |  |
| --- | --- | --- | --- | --- |
| **Antibody** | **Clone** | **Manufacturer** | **Cat#** | **Dilution** |
| CD244 | 2B4 | BD bioscience | #553306 | 1:100 |
| CD244 | eBio244F4 | eBioscience | #17-2441-82 | 1:100 |
| CD3e | 145-2C11 | eBioscience | #11-0031-85 | 1:300 |
| CD4 | RM4-5 | BioLegend | #100540 | 1:400 |
| NK1.1 | PK136 | eBioscience | #12-5941-83 | 1:200 |
| CD8a | 19517 | eBioscience | #11-0081-82 | 1:300 |
| CD19 | 1D3 | BD bioscience | #551001 | 1:500 |
| CD45 | 30-F11 | BioLegend | #103116 | 1:500 |
| CD45 | 30-F11 | BioLegend | #103128 | 1:500 |
| CD45 | 30-F11 | BioLegend | #103114 | 1:500 |
| F4/80 | BM8 | eBioscience | #17-4801-82 | 1:200 |
| F4/80 | BM8 | eBioscience | #11-4801-82 | 1:200 |
| CD11b | M1/70 | eBioscience | #56-0112-82 | 1:500 |
| CD11b | M1/70 | BioLegend | #101226 | 1:250 |
| CD11c | N418 | BioLegend | #117310 | 1:500 |
| CD11c | N418 | BioLegend | #117324 | 1:250 |
| MHCII | M5/114.15.2 | BioLegend | #107630 | 1:500 |
| Ly6G | 1A8 | BioLegend | #127618 | 1:400 |
| Ly6G | 1A8 | BioLegend | #127622 | 1:200 |
| Ly6C | HK1.4 | BioLegend | #128006 | 1:200 |
| Ly6C | HK1.4 | BioLegend | #128024 | 1:200 |
| CD206 | C068C2 | BioLegend | #141708 | 1:200 |
| IFN-γ | XMG-1 | BD bioscience | #554412 | 1:200 |
| IFN-γ | XMG-1 | BD bioscience | #560660 | 1:200 |
| Granzyme B | NGZB | eBioscience | #11-8898-82 | 1:50 |
| CD44 | IM7 | BD bioscience | #560570 | 1:400 |
| CD62L | MEL-14 | BioLegend | #104412 | 1:200 |
| PD-1 | 29F.1A12 | BioLegend | #135210 | 1:200 |
| TIGIT | 1G9 | BioLegend | #142106 | 1:100 |
| CD80 | 16-10A1 | BioLegend | #104708 | 1:200 |
| Fixable Viability 620 |  | BD bioscience | #564996 | 1:200 |
| Fixable Viability 700 |  | BD bioscience | #564997 | 1:1000 |
| Zombie NIR™ Fixable Viability Kit |  | BioLegend | #423106 | 1:1000 |

